# Effect of MIND Diet Intervention on Cognitive Performance and Brain Structure in Healthy Obese Women: A Randomized Controlled Trial

**DOI:** 10.1101/2020.04.28.065813

**Authors:** Golnaz Arjmand, Mojtaba Abbas-Zadeh, Mohammad Hassan Eftekhari

## Abstract

**Background and Aim:** Previous studies suggested adherence to recently developed Mediterranean-DASH Intervention for Neurodegenerative Delay (MIND) associated with cognitive performance. There was no prior Randomized controlled Trial (RCT) to investigate this association. This study aimed to examine the effect of MIND dietary pattern on features of cognitive performance and also changes in brain structure in healthy obese women.

**Methods:** As a total of 50 obese women assessed for eligibility, we randomly allocated 40 participants with mean BMI 32 ± 4.31 and mean age 48 ± 5.38 years to either calorie-restricted modified MIND diet or a calorie-restricted standard control diet. Change in cognitive performance was the primary outcome measured with a comprehensive neuropsychological test battery. We also performed voxel-based morphometry as a secondary outcome to quantify the differences in brain structure. All of the measurements administered at baseline and three months follow up.

**Results:** Thirty-seven participants (MIND group=22 and control group=15) completed the study. The results found in the MIND diet group working memory +1.37 (95% CI: 0.79,1.95), verbal recognition memory +4.85 (95% CI: 3.30,6.40), and attention +3.75 (95% CI: 2.43,5.07) improved more compared with the control group (*ps* < 0.05). Results of brain MRI consists of an increase in surface area of inferior frontal gyrus in the MIND diet group. Furthermore, the results showed a decrease in the cerebellum-white matter and cerebellum-cortex in two groups of study. Still, the effect in the MIND group was greater than the control group.

**Conclusions:** The study findings declare for the first time that the MIND diet intervention can reverse the destructive effects of obesity on cognition and brain structure, which could be strengthened by a modest calorie restriction.

## 1. Introduction

Obesity, characterized by excess accumulation of fat mass, is considered as one of the most growing health issues facing the world (1). Aside from the known metabolic and physiological concomitant medical risks, research supports the view that higher body mass index (BMI) is associated with alteration in global cognitive performance (2) as well as overall brain volume (3, 4). Results of a longitudinal cross-sectional study in over 2000 middle-aged adults revealed that overweight and obese people recall fewer words and took a long time to complete the cognitive tests in comparison with normal-weight participants (5). Similar results observed in a wide age range (20-82 years) research on obesity and overweight individuals illustrated that obesity without interaction with aging also has a devastating effect on cognitive performance (6).

However, it is not precisely clear how obesity can affect cognitive performance and brain structure. It has been found that negative consequences of obesity itself, before the occurrence of the related diseases can lead to cognitive impairments (4). These deficits observed by using a 5-day high-fat diet (75% energy) in healthy young men, showing that this duration was sufficient to reduce the speed of retrieval and attention. (7). Specifically, excessive energy intake and obesity go with chronically elevated levels of pro-inflammatory factors can impair the ability of neurons to adapt to chronic inflammation (8) and oxidative stress (9).

Although, only a handful of experimentally controlled studies measured the effect of diet on cognition and brain volume. Observations in human studies provided emerging insight into modifiable lifestyle factors, such as dietary patterns (10). In particular, evidence from epidemiologic studies suggested that the Mediterranean and Dietary Approach to Stop Hypertension (DASH) diet can have protective effects on cognitive decline, but results are inconsistent (11, 12). A prospective population-based study included 3831 men and women >65 years old revealed that higher DASH and Mediterranean diet score is associated with a higher average of Modified Mini-Mental State Examination Survey test (MMSE) (13). Nonetheless, neither of these two dietary patterns have been explicitly raised designed for brain health.

In 2015, Morris and colleagues developed a new brain-protection pattern. This diet has been designed after the Mediterranean and DASH diet to improve some of their dietary factors and have the highest impact on brain health and cognitive performance (14). The MIND diet emphasizes the consumption of fruits in particular berries, green leafy vegetables, nuts, olive oil, whole grains, fish, beans, poultry. The MIND diet also limits the consumption of butter, cheese, red meat, fried foods, and sweets. To investigate whether MIND diet can slow cognitive decline with aging, Morris et al. examined 960 participants over an average of 4.7 years to show that the MIND diet slows the process of reducing age-related cognitive abilities, including episodic memory, semantic memory, and the speed of perception of concepts.

Additionally, a higher score for the MIND dietary pattern is associated with a reduction in the rate of cognitive dysfunction in healthy older adults (14, 15). It seems that increased sensitivity to long-term effects of oxidative stress and inflammation due to obesity on the nervous system can reduce the cognitive and motor function of the brain. Therefore, a MIND diet with high levels containing polyphenols and antioxidant components can reverse the mechanism of oxidative stress and inflammation.

It is predicted that obesity in middle age not only associated with an increased risk of dementia in later life but also with midlife deficits in domains of language, motor function, memory performance, and most consistently implicated in executive function. These findings suggest that obesity may affect cognition before any cognitive decline appeared. Therefore, if obesity can destroy cognition in midlife, obesity-related intervention may improve this defective cycle (16).

To the best of our knowledge, the majority of the previous MIND diet studies have been cross-sectional and longitudinal, focusing on elderly participants. Here, we designed a randomized controlled trial to address this question of whether MIND dietary pattern can improve cognitive performance in middle-aged obese individuals. For this purpose, we assessed the changes in cognitive performance and the brain structures in healthy obese women.

## 2. Material and Methods

### 2.1. Participants and Sample size

This current randomized parallel well-controlled trial was carried out on 37 randomly selected participants signed up through public advertising at Imam Reza clinic of Shiraz University of Medical Sciences. To calculate sample size, we use the formula suggested for randomized clinical trials. We considered the type I error of 5% (α = 0.05), type II error of 20% (β = 0.02, power = 80%), and working memory capacity as a primary outcome variable (17). Based on this, we reached a sample size of 11 women in each group. Taking into account an estimated drop-out rate, we increased the samples.

### 2.2. Inclusion and Exclusion criteria

The inclusion criteria were defined as middle-aged women (40-60 years), without any metabolic complication, BMI 30-35 kg/m^2^, MMSE ≥ 24, and no history of severe untreated medical, neurological, and psychiatric diseases which may interfere with the study intervention. We also included participants who did not have gastrointestinal problems, did not participate in weight loss programs, or did not use weight loss drugs in the last three months. In this present study, we excluded those participants who had not wholly follow the dietary pattern or became pregnant and undergo special medical treatments during three months follow up. The eligibility of participants was first asked during a telephone screening.

### 2.3. Procedure

Participants were randomly allocated to the calorie-restricted control diet and calorie-restricted modified MIND diet group using a computerized, web-based random number table. The same dietitian who was blinded generated the random allocation sequence and assigned participants to each group. At the initial visit, participants underwent a face to face standardized medical interview and also a neurological examination. Demographic and anthropometric data, as well as details on dietary intakes, have been gathered. Nutritional data were collected using a 168-item semi-quantitative food frequency questionnaire (FFQ) to know the usual dietary intake of participants (18). Before and after the three months of study, participants underwent a comprehensive neuropsychological test as well as structural neuroimaging. All study protocols were approved by the Ethics Committee of Shiraz University of Medical Science, and participants were explained the ethical aspect of the study. Participants also provided signed informed consent before participation following the Declaration of Helsinki Law (IR.SUMS.REC.1397.759).

### 2.4. Diets

The MIND dietary pattern used for this study was based on the MIND diet developed by Morris and colleagues in 2015 (14). Participants in the MIND diet group received instruction in modifying the content of their diet to meet MIND pattern guidelines. This pattern emphasizes natural, plant-based foods, specifically promoting an increase in consumption of berries and green leafy vegetables, whole grain cereals, fish, nuts, and olive oil, with limited intakes of animal-based and high saturated fat foods. The component and scoring of the MIND diet were shown in table 1. In the current study, wine consumption was not included because its usage is forbidden in our country. Therefore, we encourage our participants to use grape and grape juice, as well as raisins and currant to modify servings.

**Table 1.** MIND diet components and scoring

| MIND diet components | Score |  |  |
| --- | --- | --- | --- |
|  | 0 | 0.5 | 1 |
| **Green leafy vegetables<sup>a</sup>** | ≤ 2 serving/week | >2 to < 6/week | ≥2 serving/week |
| **Other vegetables<sup>b</sup>** | ≤ 5 serving/week | 5 to < 7/week | ≥ 1 serving/day |
| **Berries** | ≤ 1 serving/week | 1-2/week | ≥ 2servings/week |
| **Nuts** | < 1/month | 1/month to <5/week | ≥ 5 servings/week |
| **Olive oil** | Not primary oil | - | primary oil |
| **Butter, margarine** | > 2 table spoon/day | 1-2 table spoon/day | < 1 table spoon/day |
| **Cheese** | ≥ 7 servings/week | ≥ 1-< 7/week | < 1 servings/week |
| **Whole grains** | < 1 serving/day | ≥ 1-<3/day | ≥ 3 servings/day |
| **Fish (not fried)** | Rarely | 1-3/month | ≥ 1 meals/week |
| **Beans<sup>c</sup>** | < 1 meal/week | 1-3/week | > 3meal/week |
| **Poultry (not fried)** | < 1 meal/week | ≥ 1-<2/week | ≥ 2 meals/week |
| **Red meat and products** | > 6 meals/week | ≥ 4-≤ 6/week | < 4 meals/week |
| **Fast fried foods** | ≥ 4 times/week | 1-< 4/week | < 1 time/week |
| **Pastries and sweets** | ≥ 7 servings/week | ≥ 5-< 7/week | < 5 servings/week |
| **Wine** | >1 glass/day or never | 1/month to 6/week | 1 glass/day |
| **Total score** | 0 | 7.5 | 15 |

<sup>a</sup>Kale, collards, greens; spinach; lettuce/tossed salad. <sup>b</sup>Green/red peppers, squash, cooked carrots, raw carrots,
broccoli, celery, potatoes, peas or lima beans, tomatoes, tomato sauce, string beans, beets, corn, zucchini/summer
squash/eggplant, coleslaw, potato salad. <sup>c</sup>Beans, lentils, soybeans.

A 7-day menu was then developed, meeting the required number of servings per day every week. The participants did not receive any financial compensation or gifts, but participants had to adopt the recommended diet strictly by themselves after intensive instruction and education. Participants were followed up by the same dietitian every week. Dietary intervention adherence was measured via two means of assessment, 1) MIND diet score questionnaire and 3-day food recall. Each recorded meal and snack were rated as either following the MIND diet or not. Participants were classified as having successfully adhered to the MIND diet if 80% or more of their meals/snacks met this criterion. In both groups of study, we restricted calorie intake to at least 1500 kcal/d, in which the recommended daily calorie intake was individually specified based on the World Health Organization (WHO) formula for overweight and obese women > 19 years and older (19). Meals in both calorie-restricted diet groups included a balanced mix of foods with 50-55% carbohydrates, 30% fat, and 15-20% proteins. Participants in the control group were instructed to calorie-restricted diet alone during the period of study. The control group also received general oral and written information about healthy food choices due to ethical concerns and to keep participants in the study. All parameters were collected at baseline and the end of three months of intervention.

### 2.5. Primary endpoint

#### 2.5.1. Assessment of cognitive performance

All of the participants were tested on a complex battery of neurocognitive tests to examine their cognitive performance in different domains. Verbal short memory composite included the score from Forward Digit Span Task (FDST) and Backward Digit Span Task (BDST). The working memory capacity was measured by the Letter Number Sequencing Task (LNST). Attention and visual scanning were obtained by the Symbol Digit Modality Task (SDMT). Auditory Verbal Learning Test (AVLT) was used to test verbal recognition memory performance (20). Finally, the Trail making test A and B were performed to measure executive function and task switching. We also assessed the ability to inhibit cognitive interference by well-known Stroop task.

### 2.6. Secondary endpoint

#### 2.6.1. Clinical and anthropometric data

Body weight was measured using a digital scale to the nearest 0.1 kg, and standing height was measured using a stadiometer to the nearest 0.1 cm. Body composition was obtained with a Bioelectrical Impedance Analysis (BIA) device according to a standardized protocol (In Body S-10, USA). Participants were instructed not to participate in extreme physical activity or to consume alcohol and caffeinated beverages within 24 hours before the measurement.

#### 2.6.2. Image acquisition and processing

Magnetic resonance imaging was carried out by a 3T Siemens Skyra system with a 12-channel head coil on a subgroup of 11 participants in each group. Pre- and post-scans involved high resolution T1-weighted anatomical images. T1-weighted images consisted of 192 slices and were used in the sagittal plane, which prepared with gradient-echo sequence (repetition time = 1900ms, echo time = 2.52ms, flip angle = 9°, voxel size = 1×1×mm). Image preprocessing was performed by the Freesurfer, stable version 5.1 (http://surfer.nmr.mgh.harvard.edu).

To capture the changes in brain structure in response to MIND diet intervention in study groups and reduce biased analysis and measurement noise, the longitudinal processing procedure of Freesurfer was used. For this procedure, first, the images from baseline and three months were independently processed. Then, the baseline data was subtracted from the three months data in a vertex-wise manner. For group analysis, the output map for each participant was resampled, normalized to common space, and smoothed with a Full Width of Half Maximum (FWHM) of 10 mm. Smoothing was only performed for vertex-wise whole-brain analyses, and it was not used for parcellation measures of cortical regions and volumetric measures of subcortical regions.

### 2.7. Statistical analysis

Data were analyzed using SPSS 22.0 for Windows (SPSS, Chicago, IL) and level of significance for all analyses adjusted at alpha *p* < 0.05. To screen the normality of variables, a one-sample Kolmogorov-Smirnow test was used. Descriptive statistics for neuropsychological data and physical parameters are performed as mean ± SD. Independent sample t-tests were used to determine the differences between baseline measures of two groups. A paired sample t-test was used to compare the data at the baseline and the three months of intervention for each group. We also used the Man-Whitney as a non-parametric test to compare the differences in brain structure between the two groups. A mixed model two-way repeated-measure analysis of variance (ANOVA) was performed to compare the mean differences between groups that had been divided into two factors, where time (baseline, three months) as a within-participant factor and treatment (MIND diet group, control group) as a between-participant factor. The effect of diet intervention in the whole brain and Regions of Interest (ROIs) was statistically tested. For whole-brain analysis, output maps were corrected for multiple comparisons using the false discovery rate (FDR) 0.05 as implemented in Freesurfer. Corresponding effect sizes in the form of partial Eta square were calculated to evaluate the magnitude of effect for all hypotheses.

## 3. Results

### 3.1. Participant characteristics

The study took place between October 2018 and March 2019. We screened 50 volunteers for inclusion criteria, and 40 people met the inclusion criteria. The analyzed study consisted of 37 included participants (n = 22 MIND diet group, n = 15 control group). The CONSORT flowchart diagram has been reported in figure 1. The participants in both groups completed the study, and no side effects have been reported. Descriptive results showed that at baseline, the mean (SD) age of participants was 48 ± 5.3 years, and the majority of them were married (83.8%). All of the participants who participated in the study were right-handed (21). Table 2 shows the baseline characteristics of the participants who participated in the study. The MIND diet and control groups had similar clinical characteristics, and no participants reported a history of diabetes and hypertension. To examine of anthropometric parameters over three months, a repeated measure ANOVA analysis using the variables as within-subject factor and the type of treatments as a between-subject factor was performed. Figure 2 shows that significant group × time interaction for the body weight and percent of body fat after a three months intervention in the MIND diet group in comparison with the control group (*ps* < 0.05, fig 2. Panel A and B).

**Table 2.** Baseline characteristics of participants according to the group studies.

| Variables | MIND diet<br>(n:22) | Control group<br>(n:15) | P-value |
| --- | --- | --- | --- |
| Age (years) | 48.95 (5.03) * | 48.86 (6.04) | 0.962 |
| Education (years) | 16.40 (1.05) | 16.40 (1.05) | 0.980 |
| Weight (kg) | 81.95 (10.96) | 82.33 (15.35) | 0.930 |
| Height (cm) | 160.18 (4.65) | 159.60 (5.50) | 0.731 |
| Percent of body fat (%) | 40.84 (5.34) | 41.05 (6.32) | 0.913 |
| Fat Free Mass (kg) | 48.05 (5.14) | 47.66 (5.33) | 0.823 |
| BMI (kg/m<sup>2</sup>) | 31.90 (3.70) | 32.19 (4.99) | 0.839 |
| WC (cm) | 99.75 (9.78) | 103.69 (12.61) | 0.292 |
| MMSE (score) | 26.22 (1.63) | 26.73 (1.75) | 0.375 |

\* mean (SD)= Baseline differences between groups used t-test analysis.
Abbreviations: BMI = Body Mass Index, WC = Waist circumference, MMSE = Mini-Mental State Examination survey.

**Figure 1.**
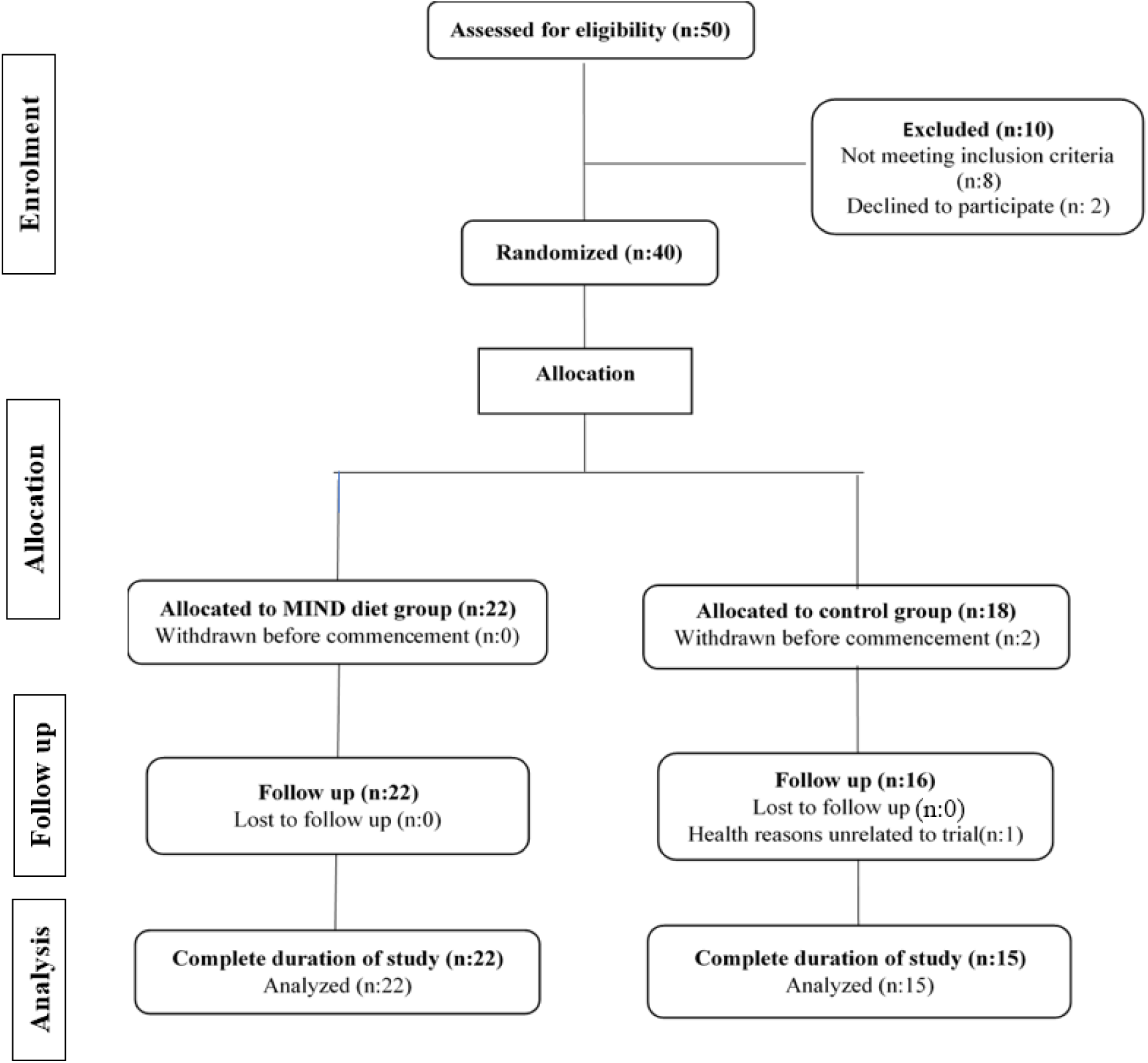
CONSORT flow diagram of the study.

**Figure 2.**
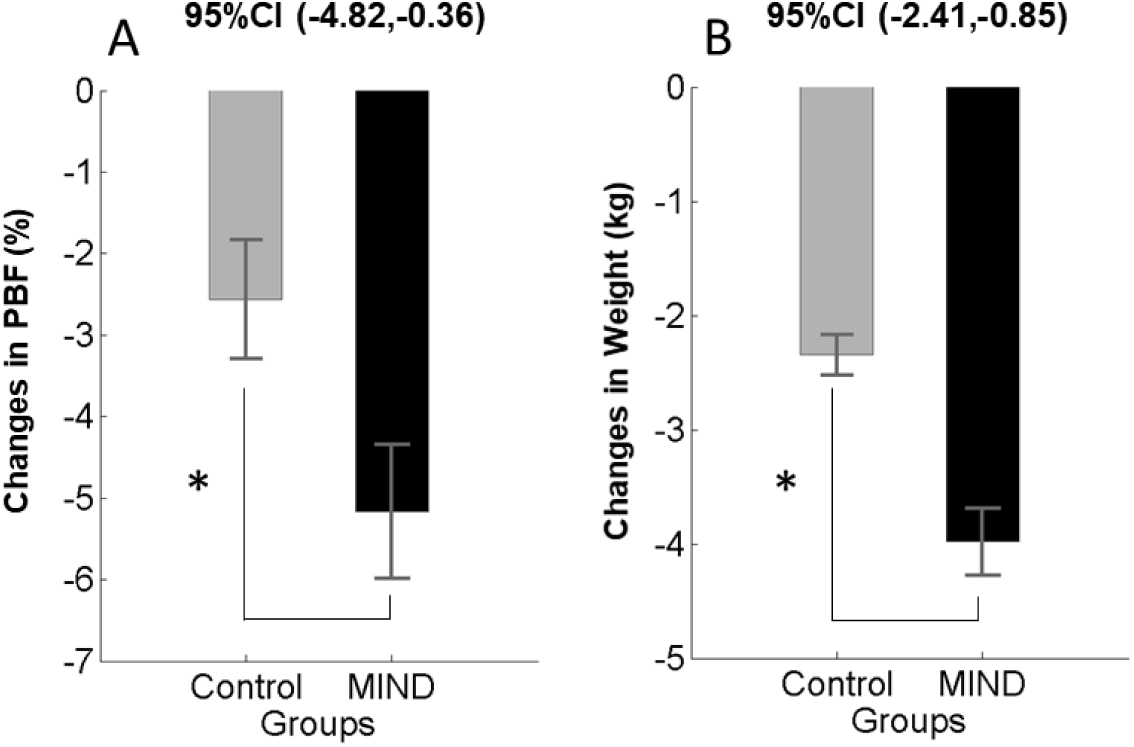
Anthropometric changes (mean and standard error of the mean) in the MIND diet group (black color) and control group (gray color) at baseline and follow up. Note that *P*-value < 0.05 in a repeated measure ANOVA test indicating significant improvement in weight (panel A) and percent of body fat (panel B) in the MIND diet group in comparison with the control group. Abbreviation: PBF= Percent of Body.

### 3.2. Changes in cognitive performance

By using linear mixed model analysis, we found that there was a statistically significant effect of time on MIND diet group induced cognitive test score of FDST, BDST, LNST, and SDMT (*p*s < 0.05), indicated the rate of these changes over time was not similar in both groups (Table 3). However, no significant group × time interaction could be found in Trail making test B (*p* = 0.161 fig 3. Panel F) and Stroop test (*p* = 0.128 fig 3. Panel G) but, the three months intervention had a statistically significant effect on Trial making test A (*p* = 0.002 fig 3. Panel E). A repeated measure ANOVA with a Greenhouse-Geisser correction determined that mean Auditory verbal learning task differed significantly between two-time points (*p* = 0<0.001 figs 3. Panel H). The within-group comparison revealed that improvements in all cognitive tests were detected in both the MIND and the control group, which could be part of the results of the learning effect and being familiar with the content of tests (table 3). Differences in each group’s cognitive performance tests at each time point were presented in figure 3.

**Table 3.** Cognitive performance data outcomes from group × time interaction after three months follow up study in the MIND diet group versus the control group.

| Outcome<br>Cognitive<br>performance | Mean<br>(SD) |  | F (1,35)<sup>a</sup> | p-value for repeated measure<br>ANOVA<sup>b</sup> |  |  | Effect size<br>(partial eta squared)<sup>c</sup> |  |  |
| --- | --- | --- | --- | --- | --- | --- | --- | --- | --- |
|  | MIND<br>(n:22) | Control<br>(n:15) | Interaction<br>term | Between<br>group | Within<br>group | Interaction<br>term | Between<br>group | Within<br>group | Interaction<br>term |
| **FDST** |  |  |  |  |  |  |  |  |  |
| Baseline | 7.09 (1.92) | 6.86 (1.59) | 35.515 | 0.039 | $\leq 0.001$ | $\leq \mathbf{0.001}$ | 0.116 | 0.659 | 0.504 |
| 3 months | 9.18 (1.36) | 7.20 (1.37) |  |  |  |  |  |  |  |
| **BDST** |  |  |  |  |  |  |  |  |  |
| Baseline | 4.90 (1.47) | 4.26 (1.16) | 4.509 | 0.034 | $\leq 0.001$ | $\mathbf{0.041}$ | 0.122 | 0.555 | 0.114 |
| 3 months | 5.81 (1.18) | 4.73 (0.79) |  |  |  |  |  |  |  |
| **LNST** |  |  |  |  |  |  |  |  |  |
| Baseline | 6.90 (2.04) | 5.00 (1.41) | 23.101 | $\leq 0.001$ | $\leq 0.001$ | $\leq \mathbf{0.001}$ | 0.421 | 0.623 | 0.398 |
| 3 months | 8.61 (1.39) | 5.40 (1.24) |  |  |  |  |  |  |  |
| **SDMT** |  |  |  |  |  |  |  |  |  |
| Baseline | 42.68 (7.74) | 36.26 (11.06) | 33.231 | 0.008 | $\leq 0.001$ | $\leq \mathbf{0.001}$ | 0.184 | 0.700 | 0.487 |
| 3 months | 47.50 (7.16) | 37.33 (10.23) |  |  |  |  |  |  |  |
| **TMTA** |  |  |  |  |  |  |  |  |  |
| Baseline | 39.90 (12.12) | 37.00 (9.57) | 11.070 | 0.984 | $\leq 0.001$ | $\mathbf{0.002}$ | 0.000 | 0.309 | 0.240 |
| 3 months | 33.68 (7.37) | 36.46 (9.31) |  |  |  |  |  |  |  |
| **TMTB** |  |  |  |  |  |  |  |  |  |
| Baseline | 88.50 (30.35) | 101.53 (49.91) | 2.048 | 0.256 | $\leq 0.001$ | 0.161 | 0.035 | 0.327 | 0.055 |
| 3 months | 83.40 (27.54) | 99.06 (46.50) |  |  |  |  |  |  |  |
| **AVLT** |  |  |  |  |  |  |  |  |  |
| Baseline | 5.81 (2.06) | 5.40 (2.02) | 10.485 | 0.067 | $\leq 0.001$ | $\leq \mathbf{0.001}$ | 0.092 | 0.662 | 0.431 |
| 3 months | 7.81 (1.81) | 5.86 (1.76) |  |  |  |  |  |  |  |
| **Stroop** |  |  |  |  |  |  |  |  |  |
| Baseline | 113.04 (43.95) | 150.86 (64.67) | 2.434 | 0.011 | $\leq 0.001$ | 0.128 | 0.171 | 0.393 | 0.065 |
| 3 months | 92.27 (29.44) | 140.33 (59.44) |  |  |  |  |  |  |  |

Mixed Model repeated measure ANOVA results of MIND diet group versus control group on cognitive performance after three months follow up. Abbreviation: MIND = Mediterranean-DASH Intervention for Neurodegenerative Delay, FDST= forward digit span task, BDST = Backward Digit Span Task, LNST = letter Number Sequencing Task, SDMT = Symbol Digit Modalities Task, TMTA = Trail Making Test A, TMTB = Trail Making Test B, AVLT = Auditory Verbal Learning Test. <sup>a</sup>=F-value (df1 for the numerator, df2 for the denominator) , degree of freedom between-subjects, within-subjects, and the interaction term, <sup>b</sup>= P-value for mixed repeated measure ANOVA, <sup>c</sup>=Partial eta squared for the ratio of variance associated with an effect.

**Figure 3.**
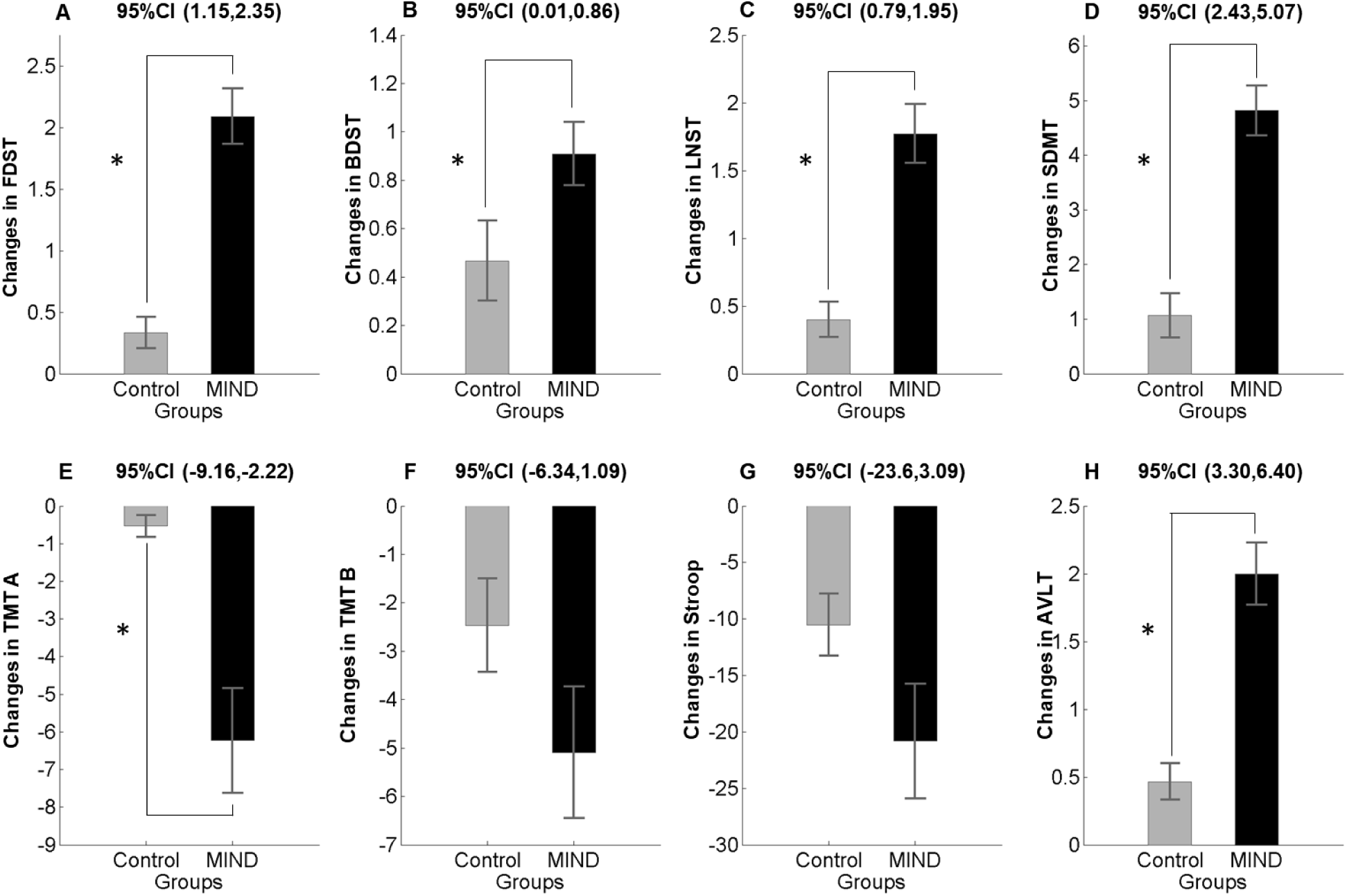
Changes in cognitive performance score (mean and standard error of the mean) in the MIND diet group (black color) and control group (gray color) at baseline and follow up. *P*-value < 0.05 in a repeated measure ANOVA determined that MIND diet intervention significantly altered the mean score of FDST, BDST, LNST, SDMT, TMTA (panel A, B, C, D, E) and AVLT (panel H). Similar but not significant trends were found for TMTB and Stroop task (panel F and G). Abbreviation: MIND = Mediterranean-DASH Intervention for Neurodegenerative Delay, FDST= forward digit span task, BDST = Backward Digit Span Task, LNST = letter Number Sequencing Task, SDMT = Symbol Digit Modalities Task, TMTA = Trail Making Test A, TMTB = Trail Making Test B, AVLT = Auditory Verbal Learning Test.

### 3.3. Change in brain structure

The whole-brain analysis did not indicate a significant effect for time × group interaction for cortical thickness, surface area, and cortical volume measures. It might be due to a few participants in each group and the short length of study. Also, we quantified the effect of diet intervention on Freesurfer generated cortical parcellation and subcortical segmentations as ROI analysis. Among cortical parcellation and subcortical segmentations, we selected areas of the brain (included: orbito-frontal cortex, inferior frontal gyrus, hippocampus, and cerebellum), which the effect of dietary patterns on them was examined previously (22, 23). To quantify the changes in brain structure between the two groups, we used the Man-Whitney u test. The result revealed significant changes in the surface area of inferior frontal gyrus in the MIND diet group (Fig 4. panel A) as the surface area in this region increase in the MIND diet group and decreased in the control group (*p* = 0.018). It was indicating that using a three months MIND diet can prevent surface area loss in the MIND group in comparison with the control group. Furthermore, the results showed a decrease in cerebellum white matter and cerebellum gray matter in two groups of studies (Fig 4. panels B & C). As can be seen in Fig 4, the effect in the MIND group was more than the control groups.

**Figure 4.**
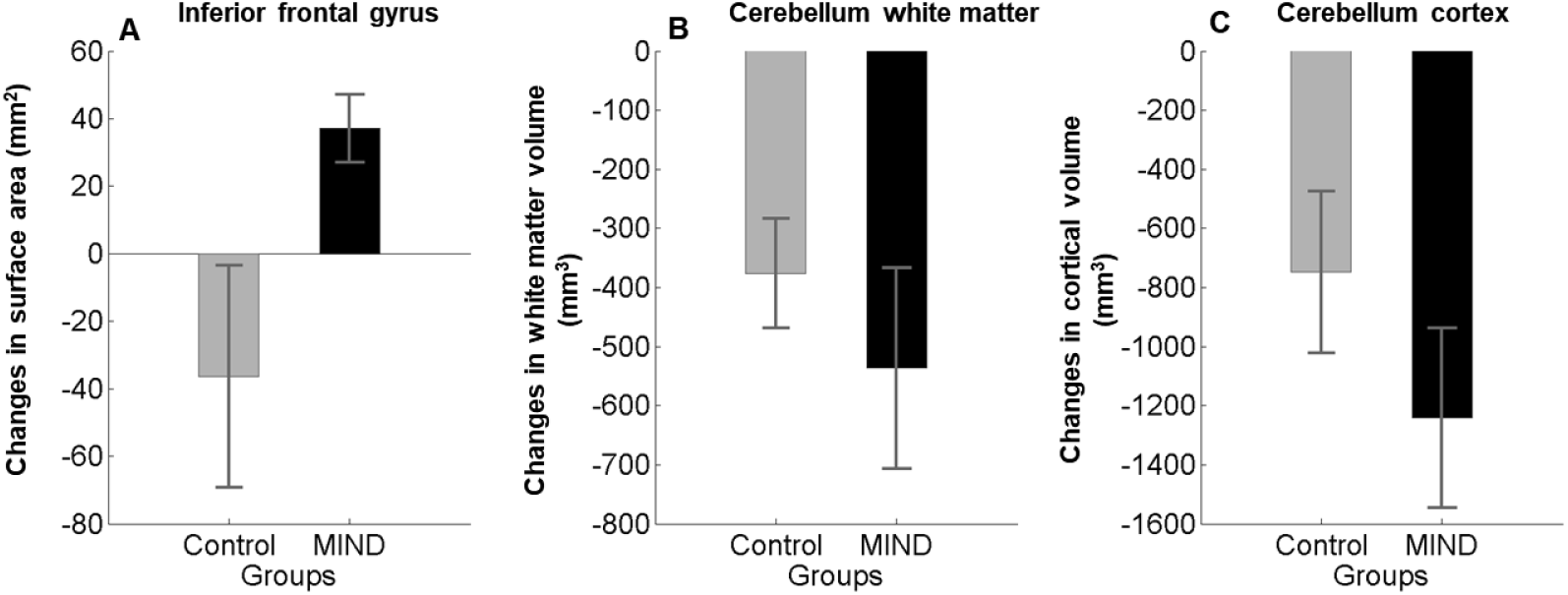
Time × group interaction for gray and white matter volumes of brain regions in the MIND diet group (black color) compared with the control group (gray color). Changes (mean and standard error of the mean) presented. Results showed that MIND diet intervention significantly increased mean changes in the surface area of inferior frontal gyrus in comparison with the control group.

## 4. Discussion

In this three-month randomized controlled study, we investigated the effect of the MIND diet on cognitive performances and brain volume changes in healthy obese women. To the best of our knowledge, this is the first study evaluating the effect of a short-term MIND diet intervention on cognition. Our results showed that the mean change of weight and percent of body fat were more decreased in the MIND diet group in comparison with the control group. Besides, group × time interaction revealed that in the MIND diet group, cognitive performance tests significantly improved relative to the control group. Our results also found that the three months intervention study can increase the volume of IFG in the MIND diet group between time points.

A distinction can be considered between MIND diet-induced weight loss and weight reduced by calorie restriction. In addition, even a moderate weight loss of about 10% or less has been shown to lead to several health benefits that improve cognitive performance. In particular, our study showed that MIND hypocaloric diet intervention had been associated with more considerable improvement in weight and percentage of body fat than the hypocaloric diet alone. In line with this thought, the current previous randomized trial on 19 obese women showed that weight loss due to calorie restriction has a beneficial effect on brain structure (24).

The results of our study indicate that adherence to the MIND diet has a more significant positive effect on FDST and BDST, which were used to measure verbal short memory. These findings in line with a meta-analysis that revealed, adherence to the Mediterranean diet significantly improved working memory and global cognition compared to controls. (25). Amelioration in cognition performance was not specific for the MIND diet group alone and could be found in the control group too. Nonetheless, this improvement was significantly greater in the MIND diet group. In addition to enhancement in working memory, we also received an improvement in Trail making test A that is considered as executive function, as well as verbal recognition memory concerning better performance on AVLT. Our findings are consistent with community based American cohort study that found a higher adherence to the Med diet was associated with a lower risk of developing MCI and slower rates of cognitive decline (26). Whereas an Australian cohort with 8-year follows up could not find an association between higher compliance of Med diet and cognition improvement (27). Additionally, Smith, Blumenthal (28) demonstrated that the DASH diet, combined with aerobic exercise and reduced-calorie, was associated with advancement in psychomotor speed performance relative to controls after the 4-month intervention. These inconsistent findings can be explained by the characteristics of the study populations, long duration of the effect of nutritional changes on cognition and the score used to evaluate adherence to the dietary patterns.

In the MIND diet group, our MRI data analysis showed an increase in gray matter volume in bilateral IFG. Our result was in line with Prehn’s study, which shows the calorie restituted diet for 12-week improved recognition memory, paralleled by an increase in gray matter volume in IFG(24). Similarly, a fMRI study of obese individuals declares that a 10 % reduction in weight loss following a low-calorie diet for 6-8 weeks was accompanied by increased activity of brain reward regions and decision-making systems such as inferior frontal gyrus and middle temporal gyrus (29). Previous human studies declare that obesity can be associated with decreasing the executive control of eating behaviors via reducing the activation of inferior frontal gyrus (30). Therefore, MIND diet intervention might be enhanced neuronal plasticity in frontal-temporal brain regions by increasing in IFG volume. Besides, our study also resulted in a considerable reduction in gray and white matter volume of the cerebellum in both MIND and control groups. A review of 16 obese participants realized that they have higher relative brain white matter volume in the cerebellum and several brain regions, which can be partially decreased by a very low-calorie diet for six weeks (31). It is investigated by fMRI studies that the cerebellum is part of the bottom-up appetite control network, which might play a particular role in some cognitive performances, including interoceptive awareness, in addition to having a role in determining energy needs and eating behavior (32). Except for an increase in the gray matter volume of IFG, there was no group by time interaction in the rest of the regions in our study results. This is possible that the smaller sample size and short duration of our study missed statistical power to detect the effect of the MIND diet on brain structure volume.

With regard to the mechanism, although the MIND diet was established on the component of the Mediterranean and DASH diets, it also has particular features that emphasize the consumption of berries, green leafy vegetables, and olive oils. As shown in previous studies, there was a linear relationship between the use of green leafy vegetables and slowing in cognitive decline. The results of this study suggest that people who consume 1 to 2 servings green leafy vegetables per day means that they are 11 years younger than those who rarely or never consume (33). In a similar way to our results, animal models’ studies were demonstrated that higher intake of berries was associated with improvement in memory and learning. These beneficial cognitive effects of berries are also repeated in the Nurse Health Study (34). Our findings are in keeping with Washington heights-Inwood Community Aging Project results, showing higher fish intake were positively associated with a larger mean cortical thickness (35). Additionally, olive oil is one of the essential key elements of the MIND diet pattern. In line with our results, a randomized trial in the sub-study of PREDIMED showed that Mediterranean intervention supplemented with extra virgin olive oil were impressive in higher cognitive scores compared with a low-fat diet among Spaniards at high cardiovascular risk (36).

On the other hand, restrictions on the intake of red meat, saturated fats, and pastries are other essential components of the MIND pattern. These components can have detrimental effects on the cardiovascular system and consequently have been related to a more considerable cognitive decline and risk of dementia (37). As a result, these nutrients may have an independent mechanism of action that synergistically protect against neurological pathogenesis. Also, it can be speculated that these components of the MIND diet could be found to protect the brain with their antioxidant and anti-inflammatory properties to protect against obesity.

The strengths of this trial included: To our knowledge, this study was the first RCT to consider the effect of MIND diet intervention on cognitive performance and brain structure among healthy obese adults. Additionally, the present study was entirely controlled intervention studies, in which all efforts to adherence to the dietary patterns are attentively considered by a well-informed nutrition consultant at the regular intervals. Finally, in the current study, we used a comprehensive cognitive test battery that had been delineated to correlate with dietary patterns in previous systematic reviews.

Our study also has limitations that must be considered when interpreting data. First, it should be noted that the short study length, along with the rather small sample size, may not allow us for further comparisons with adequate statistical power to establish between specific subgroups. However, it is consistent with sample sizes in further studies examining the effect of different dietary patterns on cognitive performance. Second, the study sample only examined a particular population and may not display the broader adult population with varying levels of education and health. However, this relative homogeneity of study participants can be attributed to the strength of the current study.

## 5. Conclusion

In conclusion, the results of this randomized controlled study have sought to consolidate the hypothesis that shows for the first time in humans a beneficial effect of MIND diet on cognition and brain structure in obese adults. In particular, the results demonstrated that these effects were specific for minimal to marked weight loss, which may have a highlighted impact on dietary patterns and cognitive performance simultaneously. According to the current development of obesity in the present century and its threatening effect on the neuronal system in adults, the strategies that focus on the reduction of stress reactivity and modulate structural functions should be viewed as more effective than pharmacological approaches. In consequence, exploring the validity of our findings in larger study samples as well as longer durations will assist researchers in developing a clear understanding of whether or not a MIND diet intervention has an evidence-informed effect on cognitive function.

## Conflict of interest statement

The authors declare no conflict of interest.

## Acknowledgments

The authors would like to thank the participants for their kind and enthusiastic corporation. This trial was financially supported by Shiraz University of Medical Science, Shiraz, Iran (Grant numbers: 97-01-84-17299).

## Statement of Authorship

Golnaz Arjmand: Conceptualization, Methodology, Data curation, Writing, Formal analysis. Mohammad Hassan Eftekhari: Conceptualization, Methodology, Review, and editing Supervising. Mojtaba Abbas-Zadeh: Methodology, Data curation, Review and editing, Formal analysis.

## Notes

### Competing Interest Statement

The authors have declared no competing interest.

## References

1. Organization WH. Adolescent obesity and related behaviours: trends and inequalities in the WHO European Region, 2002–2014: World Health Organization. Regional Office for Europe; 2017.

2. Connolly L, Coveleskie K, Kilpatrick L, Labus J, Ebrat B, Stains J, et al. Differences in brain responses between lean and obese women to a sweetened drink. Neurogastroenterology & Motility. 2013;25(7):579–e460.

3. Smith E, Hay P, Campbell L, Trollor JN. A review of the association between obesity and cognitive function across the lifespan: implications for novel approaches to prevention and treatment. Obesity reviews. 2011;12(9):740–55.

4. Debette S, Seshadri S, Beiser A, Au R, Himali J, Palumbo C, et al. Midlife vascular risk factor exposure accelerates structural brain aging and cognitive decline. Neurology. 2011;77(5):461–8.

5. Cournot M, Marquie J, Ansiau D, Martinaud C, Fonds H, Ferrieres J, et al. Relation between body mass index and cognitive function in healthy middle-aged men and women. Neurology. 2006;67(7):1208–14.

6. Gunstad J, Paul RH, Cohen RA, Tate DF, Spitznagel MB, Gordon E. Elevated body mass index is associated with executive dysfunction in otherwise healthy adults. Comprehensive psychiatry. 2007;48(1):57–61.

7. Holloway CJ, Cochlin LE, Emmanuel Y, Murray A, Codreanu I, Edwards LM, et al. A high-fat diet impairs cardiac high-energy phosphate metabolism and cognitive function in healthy human subjects. The American journal of clinical nutrition. 2011;93(4):748–55.

8. Odegaard JI, Chawla A. Pleiotropic actions of insulin resistance and inflammation in metabolic homeostasis. Science. 2013;339(6116):172–7.

9. Miller AA, Spencer SJ. Obesity and neuroinflammation: a pathway to cognitive impairment. Brain, behavior, and immunity. 2014;42:10–21.

10. van de Rest O, Berendsen AA, Haveman-Nies A, de Groot LC. Dietary patterns, cognitive decline, and dementia: a systematic review. Advances in nutrition. 2015;6(2):154–68.

11. Tangney CC, Li H, Wang Y, Barnes L, Schneider JA, Bennett DA, et al. Relation of DASH-and Mediterranean-like dietary patterns to cognitive decline in older persons. Neurology. 2014;83(16):1410–6.

12. Tangney CC. DASH and Mediterranean-type dietary patterns to maintain cognitive health. Current nutrition reports. 2014;3(1):51–61.

13. Wengreen H, Munger RG, Cutler A, Quach A, Bowles A, Corcoran C, et al. Prospective study of Dietary Approaches to Stop Hypertension–and Mediterranean-style dietary patterns and age-related cognitive change: the Cache County Study on Memory, Health and Aging. The American journal of clinical nutrition. 2013;98(5):1263–71.

14. Morris MC, Tangney CC, Wang Y, Sacks FM, Barnes LL, Bennett DA, et al. MIND diet slows cognitive decline with aging. Alzheimer’s & dementia. 2015;11(9):1015–22.

15. Morris MC, Tangney CC, Wang Y, Sacks FM, Bennett DA, Aggarwal NT. MIND diet associated with reduced incidence of Alzheimer’s disease. Alzheimer’s & Dementia. 2015;11(9):1007–14.

16. Wang C, Chan JS, Ren L, Yan JH. Obesity reduces cognitive and motor functions across the lifespan. Neural plasticity. 2016;2016.

17. McMillan L, Owen L, Kras M, Scholey A. Behavioural effects of a 10-day Mediterranean diet. Results from a pilot study evaluating mood and cognitive performance. Appetite. 2011;56(1):143–7.

18. Mirmiran P, Esfahani FH, Mehrabi Y, Hedayati M, Azizi F. Reliability and relative validity of an FFQ for nutrients in the Tehran lipid and glucose study. Public health nutrition. 2010;13(5):654–62.

19. Mahan LK, Escott-Stump S. Krause’s food, nutrition, & diet therapy: Saunders Philadelphia; 2004.

20. Psaltopoulou T, Sergentanis TN. Mediterranean diet may reduce Alzheimer’s risk. BMJ Evidence-Based Medicine. 2015;20(6):202-.

21. Oldfield RC. The assessment and analysis of handedness: the Edinburgh inventory. Neuropsychologia. 1971;9(1):97–113.

22. Prehn K, Jumpertz von Schwartzenberg R, Mai K, Zeitz U, Witte AV, Hampel D, et al. Caloric restriction in older adults—differential effects of weight loss and reduced weight on brain structure and function. Cerebral cortex. 2017;27(3):1765–78.

23. Staubo SC, Aakre JA, Vemuri P, Syrjanen JA, Mielke MM, Geda YE, et al. Mediterranean diet, micronutrients and macronutrients, and MRI measures of cortical thickness. Alzheimer’s & dementia. 2017;13(2):168–77.

24. Prehn K, Jumpertz von Schwartzenberg R, Mai K, Zeitz U, Witte AV, Hampel D, et al. Caloric Restriction in Older Adults-Differential Effects of Weight Loss and Reduced Weight on Brain Structure and Function. Cereb Cortex. 2017;27(3):1765–78.

25. Loughrey DG, Lavecchia S, Brennan S, Lawlor BA, Kelly ME. The Impact of the Mediterranean Diet on the Cognitive Functioning of Healthy Older Adults: A Systematic Review and Meta-Analysis. Advances in Nutrition. 2017;8(4):571–86.

26. Scarmeas N, Stern Y, Mayeux R, Manly JJ, Schupf N, Luchsinger JA. Mediterranean Diet and Mild Cognitive Impairment. Archives of Neurology. 2009;66(2):216–25.

27. Knight A, Bryan J, Wilson C, Hodgson JM, Davis CR, Murphy KJ. The Mediterranean Diet and Cognitive Function among Healthy Older Adults in a 6-Month Randomised Controlled Trial: The MedLey Study. Nutrients. 2016;8(9):579.

28. Smith PJ, Blumenthal JA, Babyak MA, Craighead L, Welsh-Bohmer KA, Browndyke JN, et al. Effects of the Dietary Approaches to Stop Hypertension Diet, Exercise, and Caloric Restriction on Neurocognition in Overweight Adults With High Blood Pressure. Hypertension. 2010;55(6):1331–8.

29. Rosenbaum M, Sy M, Pavlovich K, Leibel RL, Hirsch J. Leptin reverses weight loss-induced changes in regional neural activity responses to visual food stimuli. J Clin Invest. 2008;118(7):2583–91.

30. McCaffery JM, Haley AP, Sweet LH, Phelan S, Raynor HA, Del Parigi A, et al. Differential functional magnetic resonance imaging response to food pictures in successful weight-loss maintainers relative to normal-weight and obese controls. The American Journal of Clinical Nutrition. 2009;90(4):928–34.

31. Haltia LT, Viljanen A, Parkkola R, Kemppainen N, Rinne JO, Nuutila P, et al. Brain White Matter Expansion in Human Obesity and the Recovering Effect of Dieting. The Journal of Clinical Endocrinology & Metabolism. 2007;92(8):3278–84.

32. Zhu JN, Wang JJ. The cerebellum in feeding control: possible function and mechanism. Cell Mol Neurobiol. 2008;28(4):469–78.

33. Morris MC, Evans DA, Tangney CC, Bienias JL, Wilson RS. Associations of vegetable and fruit consumption with age-related cognitive change. Neurology. 2006;67(8):1370–6.

34. Devore EE, Kang JH, Breteler MMB, Grodstein F. Dietary intakes of berries and flavonoids in relation to cognitive decline. Annals of Neurology. 2012;72(1):135–43.

35. Staubo SC, Aakre JA, Vemuri P, Syrjanen JA, Mielke MM, Geda YE, et al. Mediterranean diet, micronutrients and macronutrients, and MRI measures of cortical thickness. Alzheimer’s & Dementia. 2017;13(2):168–77.

36. Martínez-Lapiscina EH, Clavero P, Toledo E, Estruch R, Salas-Salvadó J, San Julián B, et al. Mediterranean diet improves cognition: the PREDIMED-NAVARRA randomised trial. Journal of Neurology, Neurosurgery & Psychiatry. 2013;84(12):1318–25.

37. Ngabirano L, Samieri C, Feart C, Gabelle A, Artero S, Duflos C, et al. Intake of Meat, Fish, Fruits, and Vegetables and Long-Term Risk of Dementia and Alzheimer’s Disease. J Alzheimers Dis. 2019;68(2):711–22.

